# Using a substitute species to inform translocation of an endangered territorial mammal

**DOI:** 10.1101/2022.06.24.497487

**Authors:** Marina Morandini, John L. Koprowski

## Abstract

Substitute species can inform management strategies without exposing endangered species unnecessarily. Further, experimental approaches may help to identify the causes of translocation failures, leading to improve the chances of success. We used a surrogate subspecies, *Tamiasciurus fremonti fremonti* to test different translocation techniques to inform potential management actions on the endangered Mt. Graham red squirrel (*Tamiasciurus fremonti grahamensis*). We fitted VHF radio collars to 54 animals, and we monitored their survival and movements until individuals settled on a new territory. We considered the effect of season, translocation technique (soft or hard release), and body mass on survival, distance moved after release, and time to settlement of translocated animals. Survival probability averaged 0.48 after 60 days from the translocation event and was not affected by season or translocation technique. 54% of the mortality was caused by predation. Distance moved and number of days to settlement varied with season, where winter was characterized by shorter distances (average of 364 m in winter versus 1752 m in fall) and a smaller number of days (6 in winter versus 23 in fall). These data emphasize the potential of substitute species to provide valuable information on possible outcomes of management strategies for closely related endangered species.

## Introduction

Managing endangered species to ensure their long-term persistence is a central theme in modern conservation science (1,2). Endangered species are broadly characterized by small population size (3), which can lead to genetic problems such as inbreeding depression, loss of genetic variation, and accumulation of deleterious mutations (4,5), and elevated risk of extinction due to stochastic events (6). The intrinsic value of species, as well as the services and the economic benefits that biodiversity provide to humans, are key reasons to consider conservation actions to slow or reverse biodiversity losses (7).

Translocation —the intentional movement of animals for conservation purposes— has been used as a technique to mitigate the loss and depletion of endangered species (8–10). However, numerous examples worldwide have shown failures for translocation programs across a wide variety of species (9,11). Multiple factors contribute to the outcomes of translocations, and often differ among species (12). Individual characteristics (e.g. personality, physical condition, age, sex), source populations (e.g. genotype, captive vs wild born), post-release environments (e.g. predation, habitat), seasonality, and translocation techniques (e.g. handling procedures, soft vs. hard release) can all lead to varying outcomes (13). Common problems with translocations include high mortality, low breeding success, and wandering behavior associated with translocated individuals (13).

An experimental approach may help identify and mitigate factors that influence the outcome of translocations, which could reduce translocations failures (12). However, such an approach may not be feasible for endangered taxa that, ethically, should not be used for experimentation in the wild (14,15). In this context, using ecologically and genetically similar species (i.e., substitute species) to test management strategies is a valuable tool to avoid disturbance of species already facing numerous challenges (16,17). A substitute species is defined as “species or populations studied with the assumption that they show how populations of conservation concern might respond to environmental disturbance” (16). For this reason, the criteria used to choose an appropriate surrogate species is critical to obtain the most reliable information to support conservation efforts (16–18).

The endangered Mt. Graham red squirrel (*Tamiasciurus fremonti grahamensis*, (19) declined to 35 animals after the extreme decrease in population size resulting from a wildfire in 2017 (20). Given the critical status of this population, active conservation efforts that include translocation to new areas or augmentation of the population from *ex situ* bred stock should be considered. Because these actions involve translocating squirrels between areas, proper translocation techniques must be developed and evaluated. Using a surrogate species to illuminate how an endangered species may respond to conservation translocation techniques is a logical management strategy. Fremont’s red squirrel, (*T. f. fremonti*, (19)) is genetically similar to the Mt. Graham red squirrel (21). Thus, this subspecies could be a substitute species candidate to test different translocation techniques to inform management actions for the endangered Mt. Graham red squirrel.

Although assessing the outcomes of substitute species may better inform conservation decisions for other imperilled taxa, these strategies have not been thoroughly evaluated in a conservation context. In this study, we use a substitute species to examine the potential effects of seasonality, translocation technique, and body mass on the survival, post-release dispersal distances, and time to settlement of translocated animals. In particular: 1) we think body mass at the time of release, translocation type, and season will influence the survival of animals after translocation. We predict that higher survival is associated with larger animals, and those released in the fall with soft release techniques. Higher body mass indicates more fat reserves, constituting energy reservoir, (22,23) during the period when animals search for a new area to settle. Soft release allows animals to acclimate to the release area (24). Fall is characterized by the availability of a large quantity of conifer cones, the principal food source of red squirrels (25,26). 2) We predict that in winter with soft release techniques, animals will settle closer to the release site and spend less time before settling in a new site. Winter is characterized by limited food availability (27,28). In winter intraspecific competition is also lower, because most of the juveniles who did not establish a new midden likely experienced mortality, decreasing population density (29). Therefore, we expect higher site fidelity in translocated animals during winter when competition is low, and animals have more time to explore the surroundings without encountering conspecifics. Soft release, instead, allows animals to learn their new environment (24,30), therefore, to mitigate their homing response after removal of the holding pen.

## METHODS AND STUDY AREA

### Study species

We selected *Tamiasciurus fremonti fremonti* as a substitute species because it is the closest genetically and ecologically subspecies to the Mt. Graham red squirrel that is not endangered (21).

The Mt. Graham red squirrel is endemic to the Pinaleño Mountains of southeastern Arizona (31). This population is separated by desert and grasslands approximately 110 km from populations of the nearest related subspecies *T. fremonti fremonti*, located in the White Mountains. This separation occurred approximately 11,000 y ago at the end of the Wisconsin glaciation (32). Both subspecies inhabit similar forests with comparable habitat characteristics, elevations and weather conditions (33,34).

Red squirrels are diurnal mammals, active year around. Red squirrels typically have a single reproductive season focused in late spring or early summer (27,35). In western North America, red squirrels are territorial and vigorously defend the center of their territory and their larderhoard (midden) from conspecifics (27,36,37). Squirrels store conifer cones when available in their midden and in pits in the ground. Middens are necessary for survival as they provide cool, moist conditions that prevent cones from drying and opening (20), thus furnishing a reliable food supply over winter (27,28). For this reason, acquisition of a territory after natal dispersal is critical to survival and reproduction of male and female red squirrels (29,38). Forest structure around middens is important in creating a microclimate necessary for cone preservation in addition to providing nesting sites, cover and escape routes from predators, and access to foraging sites (27).

### Study Area

We studied squirrels in two study sites in the Apache-Sitgreaves National Forest in the White Mountains (Arizona, USA). The first study site was near Big Lake (UTM coordinate: 12S 647118.39216871 3750447.4959222), while the second site was near Hannagan Meadow (UTM coordinates: 12S 655714.23295814 3723357.028484). Both sites had similar elevation, between 2650 - 2750 m and mixed conifer forest type. Common species were Douglas fir (*Pseudostuga menziesii*), blue spruce (*Picea pungens*), corkbark fir (*Abies lasiocarpa*), Engelmann spruce (*Picea engelmannii*), ponderosa pine (*Pinus ponderosa*), southwestern white pine (*P. strobiformis*), and aspen (*Populus tremuloides*) (33). The sites are 32 km apart.

### Trapping and release strategies of the animals

We experimentally translocated squirrels during the fall of 2018, 2019, and 2020, and during the winter of 2018/2019 and 2019/2020. In the fall, we started translocations in August until October, and we monitored the animals until the first snow (October for 2018, November for 2019). In winter we translocated the animals from January during the first winter season, and in December during the second one, until the beginning of March. We check the translocated animals until the end of April.

We used Big Lake in fall and Hannagan Meadow in winter due to accessibility. Although it is possible season might be confounded with location, both sites have stable red squirrel populations and the same vegetation type and structure suggesting this is unlikely. We trapped squirrels with wire-mesh box live traps (Tomahawk Live Trap, Tomahawk WI: Model # 201) baited with peanuts and peanut butter. We transferred animals into a cloth handling cone (39) and marked each with unique numbered ear tag (Monel 1005-1, National Band and Tag) and coloured ear disks (1 cm Model 1842, National Band and Tag), for individual identification. We weighed each animal with a Pesola spring balance to the nearest 5g and recorded reproductive condition. Handling time was never more than 5 min to reduce stress. Before translocating an individual, each was equipped with a radio collar (SOM 2190, Wildlife Materials International, 5-7g which is less than 5% of the individual’s body mass).

During the first fall in2018, we translocated two animals 900 m from the point of capture, and each returned to their territory within a few hours. For all other translocations, we used a distance > 3000 m. All animals were translocated to areas inhabited by other red squirrels due to difficulty of finding areas that had no established squirrels yet met the environmental requirements for settlement.

We implemented the hard release technique by trapping animals and transferring each in nest boxes to the new area (34 cm H, 18cm W, 23 cm L). Nest boxes were provisioned with nesting material (hay), peanuts and peanut butter with entrance holes closed to minimize visual cues known to facilitate homing (40). We retained squirrels at their release site in their closed nest boxes during the first night at a height of 2 m.

For soft releases, we transferred individuals inside a nest box to an enclosure (152 cm H, 90cm W, 90 cm L). We provided the enclosure with approximately 400 locally collected spruce cones and a feeder with peanuts, peanuts butter, and rodent chow as supplemental feeding. We kept the squirrels in the enclosure for 5 days. After five days, we opened the enclosure and let the animals leave when they chose to do so. The sample size for soft release (n=12) is smaller than hard release (n=42) because we were able to obtain the required permit and IACUC approval only in late fall 2019.

We supplemented food of animals translocated in winter (peanuts, peanuts butter, and rodent chow) using a feeder until April. We initially provided 500 grams of peanuts, 500 g of rodent chow and 4 tablespoons of peanuts butter and checked the feeder every two weeks and we refilled when necessary.

### Telemetry

We used digital receivers (Communication Specialists Inc. R-1000 receiver) and yagi 3-element directional antennas (Wildlife Materials Inc., Murphysboro IL, USA) to track each squirrel’s movements from capture to settlement, locating all individuals a minimum of once per day during the first 5 days after translocation. Subsequently, squirrels were tracked at least once per week until settlement, death, or they were missing from the study area.

We defined animals as settled when they exhibited territorial behaviour, including the rattle vocalizations or caching of cones (29), or if they remained in a 100-m radius for at least 3 consecutive days. The first day that individuals exhibited territorial vocalization or cone caching was considered the day of settlement; when staying within the same area for at least 3 days, we considered the 3^rd^ day as the settlement day. When possible, we trapped animals after settlement to monitor the change in body mass. We defined animals as missing if their signals disappeared during the season (we checked for animals consistently through all seasons) and were never detected subsequently.

We defined 3 types of mortality: unknown predator, avian predator, unknown cause. We identified avian-caused mortalities by the presence of plucked fur, a clipped tail, gut piles, and other miscellaneous parts such as ears; raptor feces or “white wash” was also often located nearby (41). When we found only a radio-collar, we defined mortality as predation from an unknown predator. When the signal was constantly from a tree with no additional movement, but it was not possible to reach the cavity, we defined the mortality as unknown cause. When we were able to recover the body of the animal, we sent the corpse to the Arizona Veterinary Necropsy Lab, Tucson, Arizona for a detailed post-mortem analysis.

### Statistical analysis

We used Proportional Hazard Regression (42) to estimate survival for all translocated animals, and the post-settlement survival. This approach models event rates (failures) as a log-linear function of predictor variables. In our case, we used body mass (continuous variable), season and type of release (both categorical predictors). Regression coefficients give the relative effect of each covariate on the survivor function (43).

We used a generalized linear model (GLMs, (44) to examine the effect of season (fall or winter), type of release, and body mass on two response variables: time to settlement (d) and distance to settlement (m). Time to settlement was modelled using a Poisson distribution with log-link, distance of settlement was modelled using a Gaussian distribution with identity link. We used generalized mixed model (with Gaussian distribution, identity link, and individual treated as a random effect) to determine if season and translocation event influence the body mass of squirrels.

## RESULTS

In total, we translocated 54 individual red squirrels (Table 1) and monitored post-translocation for 3 falls and 2 winters (2018-2020).

**Table 1:** Total number of red squirrels, *Tamiasciurus fremonti fremonti* translocated in the White Mountains (Arizona, USA) from 2018 to 2020, by sex, season (fall and winter), and translocation type (soft and hard).

| ***SEASON*** | ***HARD*** | ***SOFT*** | ***TOTAL*** |
| --- | --- | --- | --- |
| ***FALL*** | 21 | 6 | 27 |
|  | (11 males, 10 females) | (3 males, 3 females) |  |
| ***WINTER*** | 21 | 6 | 27 |
|  | (9 males, 12 females) | (3 males, 3 females) |  |
| ***TOTAL*** | 42 | 12 | 54 |

### Fate of translocated animals

For the 54 animals translocated, 12 (22.2%) disappeared, 22 (40.7%) settled, 4 (7.4%) returned to their original territory and 16 (29.7%) died. If we do not consider the disappeared animals for which final fate was unknown, of the 42 animals translocated, 52% settled in a new territory, 10% returned to the original territory, and 38% died prior to settling in a new territory. After settlement, 6 of 22 animals were depredated, 4 in winter and 2 in fall (between 17 and 32 days after settlement). The fate of translocated squirrels was similar for both sexes and both seasons (Fig. 1).

**Figure 1:**
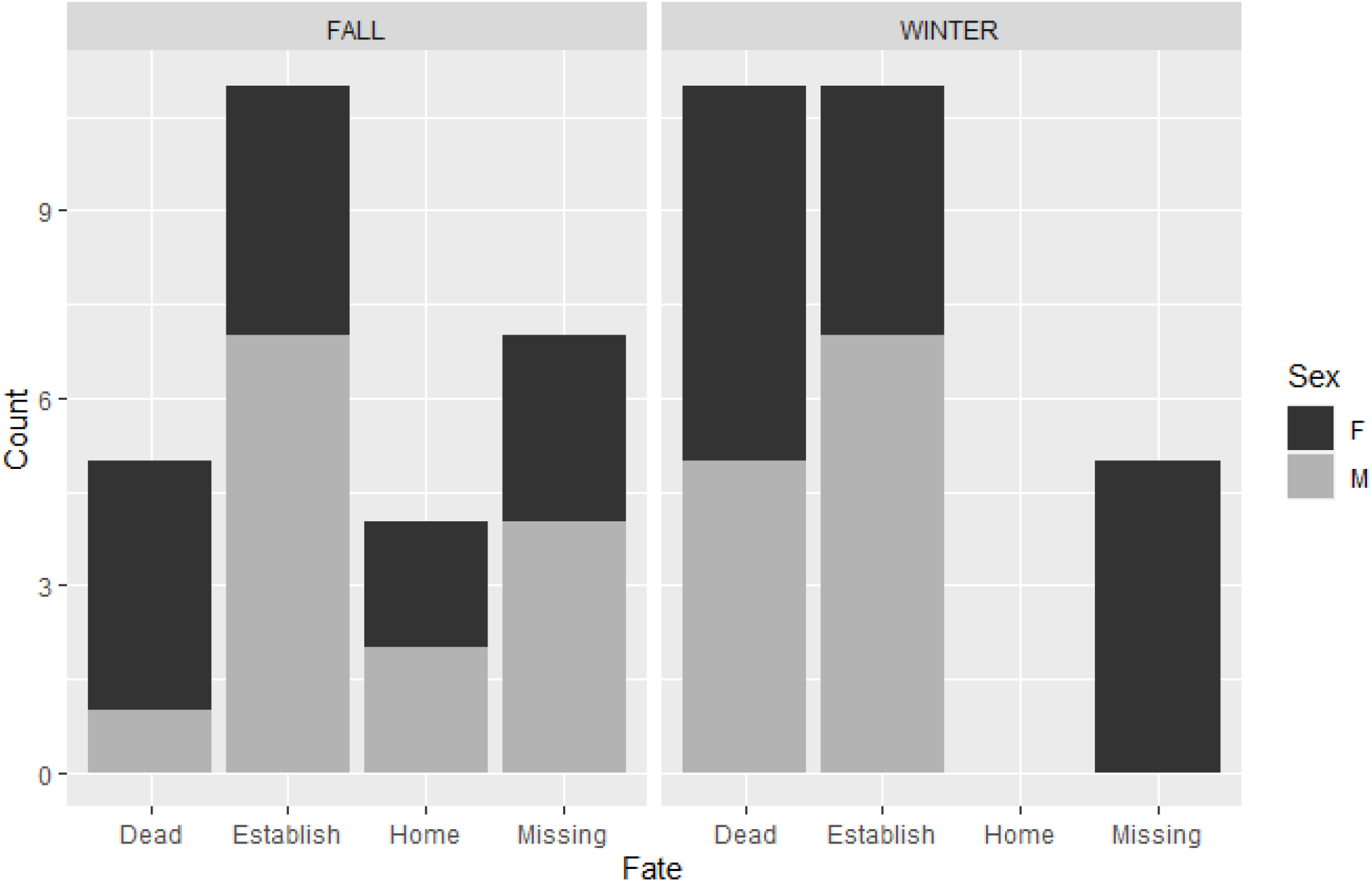
Number of red squirrels *Tamiasciurus fremonti fremonti* per different fate by sex and season (winter and fall), after translocation. The different fate has been identified as dead, establish (settled in a new area, different from the original home range), missing (when the animals was not trackable by telemetry and not possible to locate again), home (animals able to homing after being translocated).

### Mortality

In total, we confirmed 22 mortalities (40%), 15 in winter and 7 in fall. In particular, 12 (54%) died due to predation (6 by raptor and 6 by unknown predator) and 10 (46%) from unknown causes. Among the unknown causes, we had one animal with the signal in a burned area, with the collar under the snow (but impossible to recover), and 5 animals with constant signal coming from a nest, suggesting that the animals died in the nest. In fact, we were able to recover two intact squirrels from the nest where they died, one from a cavity and one from a drey. We sent, these two animals plus a squirrel found dead at the base of a tree (with intact body and no apparent signs of predation), and one found with signs of predation to a veterinary hospital for a post mortem examination. For all these animals, the pathologist considered fat stores as adequate for this species, excluding starvation as the cause of death. Except for the animal with signs of predation, which had an ear and few fingers of the front paws missing, no signs of organ damage, or wounds were recorded.

### Survival and change in body mass after translocation

Survival probability decreased steadily after translocation and stabilized below 0.5, 40 days after translocation (Fig. 2). Survival did not differ between season or translocation type; however, we observed an effect of body mass on the hazard risk, where an increase of body mass corresponded to a slightly decrease in survival (Hazard ratio 0.97, CI 0.95-0.99, p=0.039).

**Figure 2:**
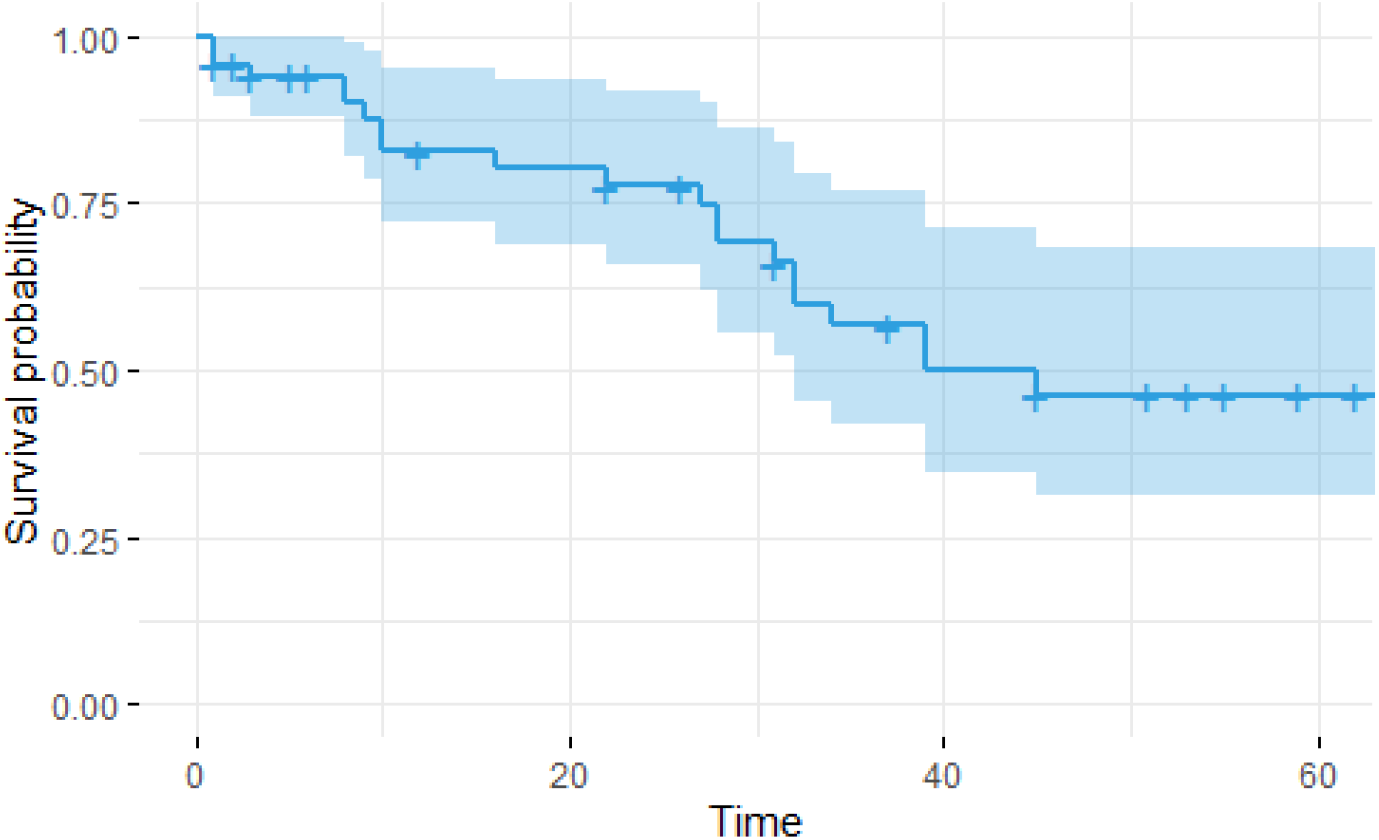
Cox Model survival curve of red squirrels *Tamiasciurus fremonti fremonti* after translocation, from the first day of release until 60 days after the release (usually corresponding also to the end of the field season). On the x-axis is represented the time in days after translocation, while on the y-axis is represented the survival probability, starting as 1 the day of the animal release and decreasing until just below 0.5 after 40 days.

We were able to document body mass for 6 of 11 (54%) animals that settled in winter and 4 of 11(36%) in fall. Translocation resulted in a decrease in animals’ body weight (t= -2.4; P =0.02), while season had no effects (t=1.04, p=0.3).

On average, we observed a loss of body mass following translocation both in winter and in fall (Fig 3). We were able to document a loss of 9.5% of body weight in an individual during only 5 days of wandering behavior (released at 280 grams and 5 days later recaptured with a weight of 255 g). After an initial loss of weight, we also recorded an increase of body mass after settlement. It was later noted that animals usually reached the same body weight prior to the translocation event in about 40 days. During this study, we documented two females with young after translocation, with the possibility that others produced offspring, but we were not able to capture to verify reproduction.

**Figure 3:**
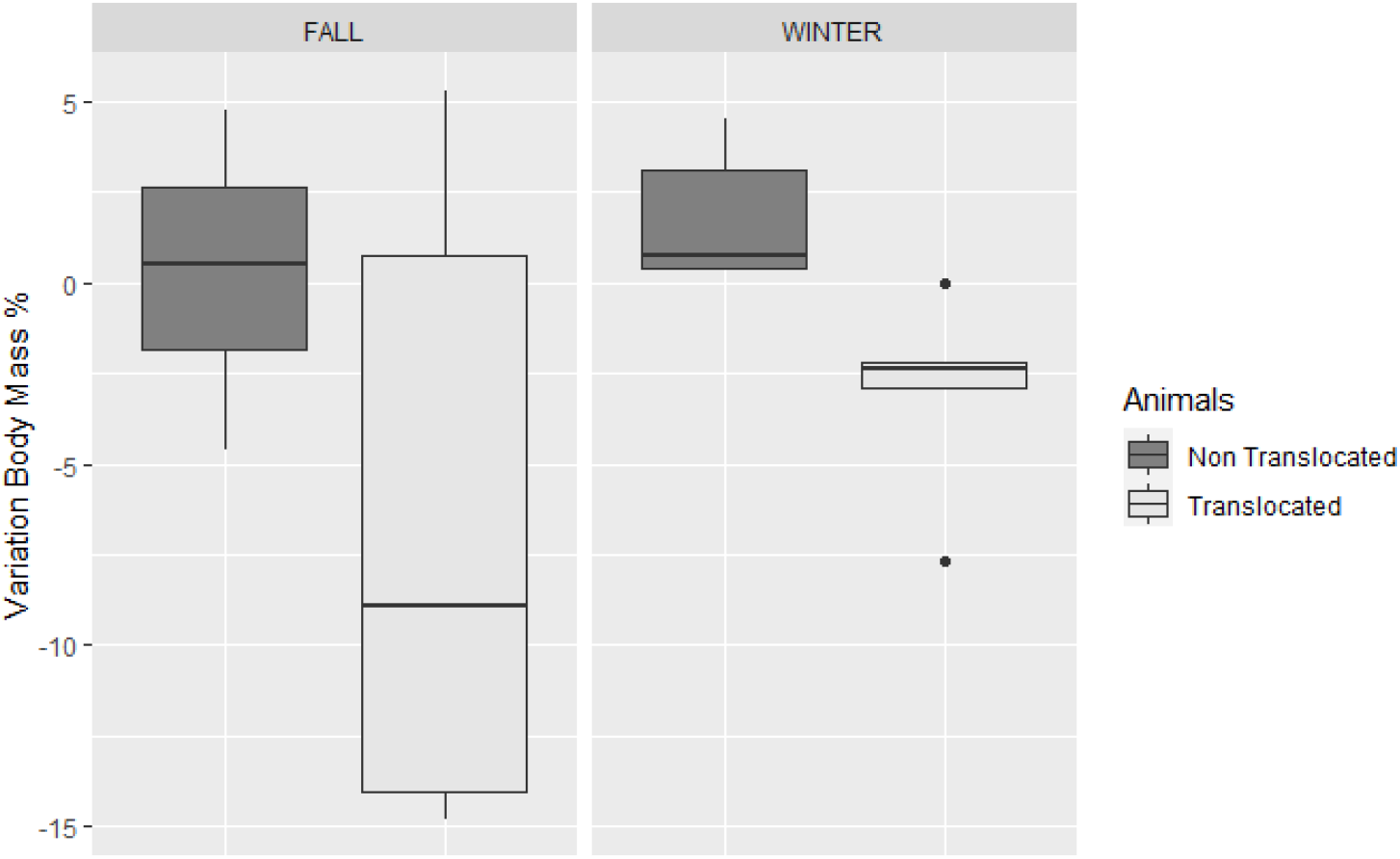
In blue the variation in body mass of translocated red squirrels *Tamiasciurus fremonti fremonti* (N=10), expressed as percentage of body mass change after translocation in respect to the weight at release. In red the variation in body mass of non-translocated animals (N=9), expressed as percentage of body mass change between two trapping sections in respect to the weight of the first trapping event (as a control group).

### Days before settlement and distance to settlement

The number of days before settlement and settlement distance were lower in winter than in fall (Fig. 4, Table 2). Body mass and type of release did not affect the number of days before settlement or settlement distance (Table 2). Intensive radio-tracking during the first week post release showed that in 20 of 48 documented cases (41%), the translocated squirrel was chased away from the release site by a resident local animal. However, in winter only 7/25 (28%) were chased away, while in fall 13/24 (54%).

**Table 2:** Estimate regression parameters, standard errors, z values and P values for the generalized linear to examine the effect of season (fall or winter), type of release, and body mass on two response variables: time to settlement (d) and distance to settlement (m) from release site.

|  | Estimate | Std. error | Z value | P value |
| --- | --- | --- | --- | --- |
| *Intercept* | 2.841 | 0.833 | 3.41 | 0.0006 |
| *Season-Winter* | -1.326 | 0.141 | -9.34 | 2 e-16 |
| *Type-Soft* | -0.111 | 0.15 | -0.73 | 0.46 |
| *Body Mass* | 0.001 | 0.003 | 0.383 | 0.70 |
| ***DISTANCE TO SETTLEMENT SITE*** |  |  |  |  |
|  | **Estimate** | **Std. error** | **t value** | **P value** |
| *Intercept* | 4153.839 | 3605.345 | 1.152 | 0.266 |
| *Season-Winter* | -1.849.538 | 527.407 | -3.507 | 0.002 |
| *Type-Soft* | -446.634 | 602.894 | -0.741 | 0.469 |
| *Body Mass* | -7.143 | 13.689 | -0.522 | 0.608 |

**Figure 4:**
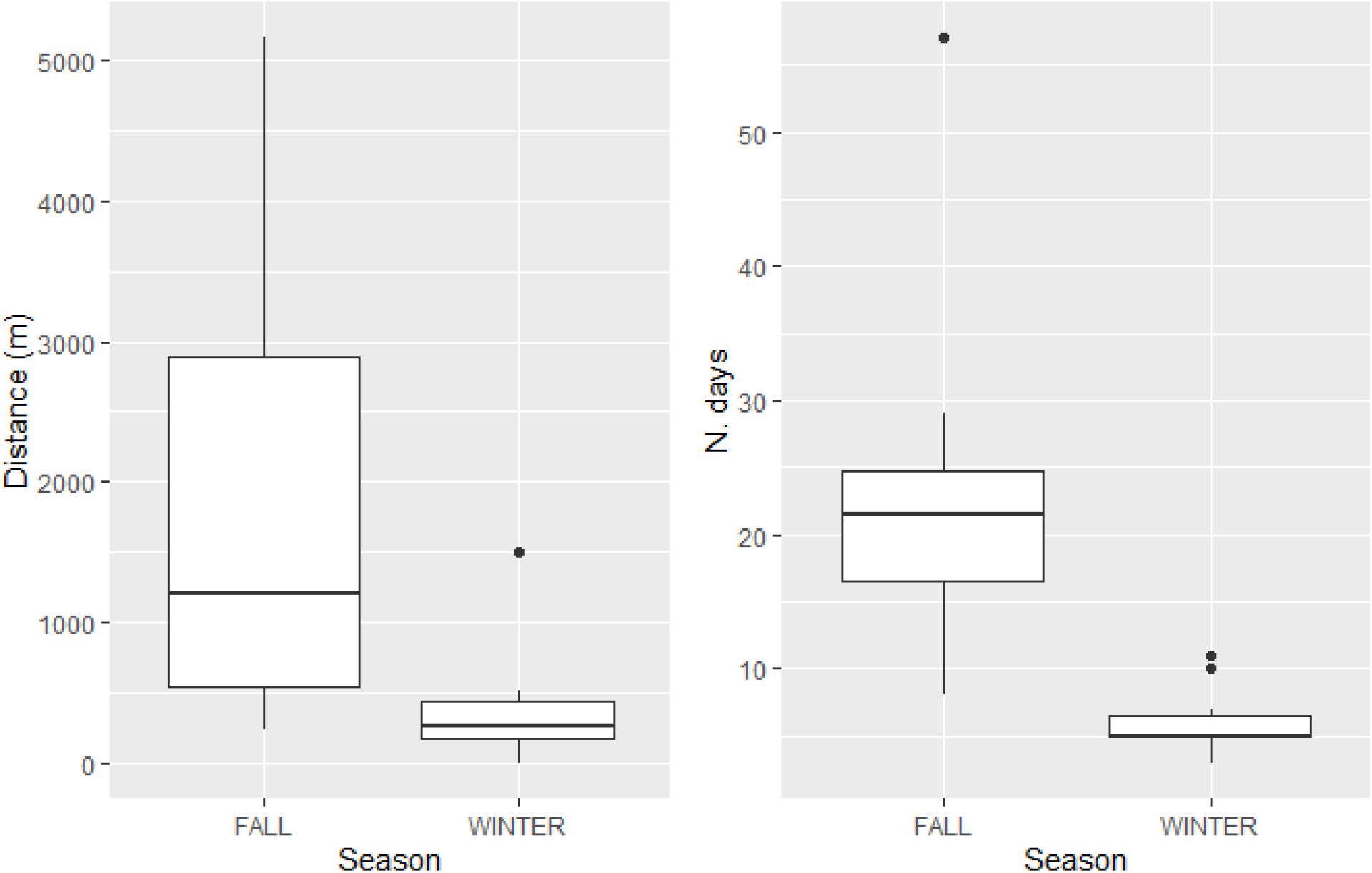
Plot of mean, sd, minimum and maximum distance of settlement site to release site (Fall: mean 1752, sd 1472, maximum 5162, minimum 233. Winter: mean 364, sd 421, maximum 1494, minimum 0 (one squirrel stayed at the release site), and number of days passed before squirrel (*Tamiasciurus fremonti fremonti*) settled (Fall: mean 23, sd 13.6, maximum 57, minimum 8. Winter: mean 6, sd 2.45, maximum 11, minimum 3).

## Discussion

We considered the effect of season, translocation technique, and body mass on survival, distance moved, and time to settlement of translocated animals. As we predicted, during winter animals showed higher site fidelity than in fall. Moreover, the distance and number of days before settlement were reduced in winter, likely due to limited food availability. Translocation techniques (soft vs. hard) did not influence days and distance to settlement site. Contrary to our prediction, season and type of release did not affect animal survival. Survival was slightly lower for animals with higher biomass, although the effect was quite small (0.97).

### Mortality after translocation: the cost of the wandering phase in a “forced dispersal”

Translocation forces animals to effectively experience dispersal. During dispersal, we observe three phases: (1) Initiation, when an individual leaves its home area; (2) Wandering, when the dispersing individual searches for new areas before settling; and (3) Settlement, when the individual settles in an area (45). Likewise, translocated animals leave the release site and wander in unfamiliar areas until they select a new place to establish. In those terms, translocation can be view as a “forced dispersal” (46). The lack of familiarity with the environment likely causes an increase of predation risk during dispersal (29), and this was similarly observed during forced dispersal after translocation. The mortality rate of animals during the wandering phase in this study was 42% (excluding the missing animals). In a forced dispersal, however, it is important to distinguish mortalities due to predation from causes connected with the translocation event (for example stress). Predation mortality accounted for 31% (excluding missing animals), a similar value than the mortality rate of red squirrels prior to settlement [22% mortality for juvenile in the forests of the Athabasca Sand Hills region of Alberta - Canada (29); 27% in the Yukon - (47); 59-67% in Rochester, Alberta - (38,48); and 23% for the subspecies, Mt Graham red squirrel - (49)].

The mortality rate in this study is also comparable to other translocated prey species. Translocated Eurasian red squirrels (*Sciurus vulgaris*) in Belgium had 50% (females) to 67% (males) mortality rates in the first months following release (50), 50% for European rabbit ((51– 53), 75% for Iberian hares (54). For each of these studies the main cause of mortality was predation, which represents a widely documented cost associated with dispersal among mammal species (55). In our study, predation accounted for at least 54% of deaths, mainly due to avian predators. Birds of prey cause 75% of deaths in juveniles and 65% in adults for the Mt Graham red squirrel in the Pinaleño Mountains for settled animals (41) as well as during dispersal (49).

Although the most common cause of mortality was predation, 10 translocated animals died from unknown causes. Through necropsy, we were able to exclude starvation for 3 of these. A tentative explanation for at least some of these unknown cases of mortality is stress, an inevitable component of translocation (56). Translocation alters stress physiology and chronic stress is potentially a major factor in translocation failure (57). Interestingly 8 of these 10 animals died during winter translocation. Even if season is not a factor that explained animal survival, mortality inside the nest was more frequent during winter than fall.

### Seasonal effects on dispersal: food availability as limiting factor in winter

Seasonality in environmental factors as well as in life-history traits can represent an important factor for the success of translocations. In resource-limited systems, different seasons are often characterized by changes in food availability and/or distribution, which in turn can influence body mass, travel distance, and mortality (58–60). We did not observe a seasonal effect on survival; however, we report an influence of season on the number of days spent by translocated individuals to settle as well as on the distance between release and settlement site. In winter, animals settled faster and closer to the release site than in fall. Food distribution probably limits the movement of the animals. In fact, the cones cached by the red squirrels in middens during the fall are fundamental to sustain the animal during winter. In the case of translocated animals during winter, the only reliable food source is the artificial food supply provided at the release site (27,28).

A second factor with potential to explain the difference between seasons on site fidelity of translocated red squirrels is intraspecific competition. In fact, habitat selection is not only affected by the suitability of the settlement site, but also by the density of conspecifics within that habitat (61). In territorial red squirrels, intraspecific competition in fall is high because juveniles disperse from their natal area and try to establish a new midden, or to take a resident’s territory (62–65). Since the behaviours associated with dispersal and competition to obtain a territory are costly, juvenile mortality between fall and the onset of winter will be high and population density will decrease (29). Therefore, in winter, animals have more time to explore the surroundings without encountering territorial conspecifics. We observed a higher number of aggressive interactions during fall than winter, with local squirrels entering the nest box of translocated animals to chase from the translocation site. Hence, our results suggest that seasonal variation in the intensity of intraspecific competition is an important factor in the settlement decisions after translocation.

### No difference between hard and soft release

We used both soft and hard releases in the translocation of red squirrels. Soft release provides translocated animals the opportunity to acclimate to their release site (30,66,67) and allows translocated animals time to learn about key aspects of their new environment, including novel stimuli (24), and potential competitors (30). However, soft release techniques do not necessarily improve results (for example higher survival) over hard release. In our study, the type of translocation (hard/soft) did not affect survival, distance to settlement site, or number of days to settlement in red squirrels. The smaller sample of soft release animals relative to hard release animals, might have affected our results. However, results from other studies also tested mixed advantages of soft release. Soft release generally improved survival, reduced movement and increased site fidelity (68). However, multiple cases exist where no difference in outcomes have been documented between hard and soft releases, as for the case of brushtail possums (69), hare-wallaby (70), and translocated colonies of black-eared miner (71).

The absence of benefit of soft release on site fidelity, can be attributed to the pressure exerted by resident conspecifics. In fact, most translocated animals (41%) were chased away by residents in the first 3 days after release. During the fall, the pressure exerted by the red squirrels present in the release area was intensified, where resident animals were exploring the enclosure and were making territorial calls from the top of the enclosure directly towards the translocated animals within the enclosure. This solitary territorial species benefits from having stable neighbours (72), at a point where animals avoid settlement in new empty middens even when they occupy poorer quality territories (73). For this reason, translocating single animals can cause the disruption of social relationship with territorial neighbours, which could have an impact on survival and behavior of translocated animals (74).

### Qualitative observations: translocation and weigh loss

We observed a loss of body mass following translocation both in winter and in fall. Many explanations are possible for this phenomenon. Translocation might cause increased physiological stress in animals, which in turn can reduce their body mass (57); or increase energetically costly behaviours, such as wandering (45). Therefore, individuals with a higher fat reserve can be better equipped to deal with low food intake during the wandering phase after translocation. In this phase animals also need to learn where to find food and shelter, and the quality of nest sites will influence energy consumption for thermoregulation, particularly when weather conditions are extreme. After an initial loss of weight, we also recorded an increase of body mass after one month post settlement. Middens are a fundamental component of red squirrels’ territory; they provide shelter, escape from predators and a reliable food supply during winter (75). For this reason, we can consider that translocated animals that settled in a new midden, successfully completed the post-translocation period. For these new residents, we expect normal behavior as well as a survival rate similar to other resident animals in the area. We observed two females in lactation, after settling in a midden, as well as males in scrotal reproductive condition. During winter, some individuals settled in an area, using artificial food, stealing cones from other middens, and moved into an existing midden only later in the season. In fall, we observed squirrels using old middens, or creating a new midden by initiating caching of cones in a new area.

### Implication for conservation management: can we use translocation on the endangered Mt Graham red squirrel?

Our study clearly documents low survival associated with translocation as well as a low settlement rates near release sites. Given these findings we do not suggest use of this conservation strategy directly on the endangered species Mt Graham red squirrel without further study. Season clearly influences the duration of spatial exploration behavior as well as the retention at the release site, mediated by food availability. In fact, during fall, large quantities of food are available in the forest (conifer seeds and mushrooms), while in winter animals need to rely on the artificially provided food supply. In fall, the most common cause of death was predation, whereas in winter most frequently animals died for unknown causes (potentially stress). The two different seasons were also characterized by different intraspecific pressure, characterized by vocalizations and frequent chases. In fact, during winter no juveniles were likely searching for a new territory. Further study to consider what the cause of death in animals during winter is necessary if this season is to be preferred, whereas in fall a means to retain animals at the release site must be identified. Perhaps the influence of neighbours in the settlement of a species could increase success. Solitary species with stable neighbours benefit from the maintenance of these social relationship during translocation (74). Red squirrels can maintain the same territory throughout their life, and consequently often choose territory sites close to familiar neighbours (63,65). Red squirrels with familiar neighbours reduced rates of territorial rattle calls and increased time spent in the nest (76). Therefore, considering the composition of the group to translocate can influence survival. Assessing a possible strategy that involves translocation of a monitored group of subjects in a defined area and moving the neighbourhood to the same translocation site to observe how translocation success changes might provide important insight. In this study, we could not translocate the species into red squirrel-free areas, therefore moving animals to areas without territorial conspecifics could potentially lead to different results due to the absence of competitors.

### Lessons from using substitute species

Our results emphasize the potential of substitute species to provide valuable information on possible outcomes of management strategies for closely related endangered species. In fact, our methodology was useful to develop and improve management strategies, including translocation, to achieve an increase in the likelihood of success (77). For example, substitute species can help to design individual marking, monitoring methods, and to test translocation techniques (78), as well as to compare soft vs hard release (79), and to refine the techniques to translocate and hand-rearing chicks (80). Substitute species also aid in the identification of key problems that can arise during phases of management plans and that may affect the related target species (Fischer & Lindenmayer, 2000). Hence, the outcomes of these trials will assist with different and better management strategies, as illustrated by the translocation of mountain lions in Florida (81).

Finally, our study shows that substitute species allow us to test management strategies, obtain an adequate sample size for statistical inference, and to test multiple scenarios, without risking the endangered species. In our case, the estimated population size of the endangered species when we planned this study (fall 2017) was only 35 individuals (20), therefore it would be impossible to test any management strategies directly on the remaining animals, without the multiple risks associated with any type of manipulation. Despite the potential offered by substitute species, not many studies used a non-endangered relative species as a substitute for an actual target species (82). Here, we have demonstrated, using a substitute species to obtain essential knowledge, how the translocation technique can be potentially detrimental when applied to an endangered species. Such results assist in delivering the appropriate methods and testing new approaches using the substitute species. We therefore conclude the value of a substitute species for testing management strategies before applying to an endangered species and suggest that such applications should be considered early in the process.

## Acknowledgment

We thank the agency partner Arizona Game and Fish Department and the University of Arizona School of Natural Resources and the Environment. We would like to thank the Mt. Graham Red Squirrel Research Program graduate and undergraduate research assistants for valuable help in the field. This research was supported by grants to JLK from the University of Arizona, Arizona Game and Fish Department (grants no. I18005 and I16002), and T & E Inc. Grants for Conservation Biology. All field work was conducted under University of Arizona Institutional Animal Care and Use Committee protocol # 16-169, Arizona Game and Fish Department scientific collecting permit # SP651773 for 2019, SP403044 for 2020, SP407072 for 2021, U.S. Fish and Wildlife Service permit # TE041875-2 and adhered to the American Society of Mammologist’s guidelines for the use of wild mammals in research (Sikes & Gannon, 2011). This manuscript was improved by comments from R. W. Mannan, L. Wauters, R. Steidl, and M.V. Mazzamuto.

## Notes

### Competing Interest Statement

The authors have declared no competing interest.

